# The vagus nerve mediates the stomach-brain coherence in rats

**DOI:** 10.1101/2022.01.25.477693

**Authors:** Jiayue Cao, Xiaokai Wang, Jiande Chen, Nanyin Zhang, Zhongming Liu

**Affiliations:** Department of Biomedical Engineering, University of Michigan, Ann Arbor, USA; Department of Electrical Engineering and Computer Science, University of Michigan, Ann Arbor, USA; Division of Gastroenterology and Hepatology, University of Michigan, Ann Arbor, USA; Department of Biomedical Engineering, Huck Institutes of the life sciences, Pennsylvania State University

## Abstract

Interactions between the brain and the stomach shape both cognitive and digestive functions. Recent human studies report spontaneous synchronization between brain activity and gastric slow waves in the resting state. However, this finding has not been replicated in any animal models. The neural pathways underlying this apparent stomach-brain synchrony is also unclear. Here, we performed functional magnetic resonance imaging while simultaneously recording body-surface gastric slow waves from anesthetized rats in the fasting vs. postprandial conditions and performed a bilateral cervical vagotomy to assess the role of the vagus nerve. The coherence between brain fMRI signals and gastric slow waves was found in a distributed “gastric network”, including subcortical and cortical regions in the sensory, motor, and limbic systems. The stomach-brain coherence was largely reduced by the bilateral vagotomy and was different between the fasting and fed states. These findings suggest that the vagus nerve mediates the spontaneous coherence between brain activity and gastric slow waves, which is likely a signature of real-time stomach-brain interactions. However, its functional significance remains to be established.

## Introduction

The stomach and the brain interact with one another. The brain regulates food ingestion and digestion (Konturek et al., 2004; Holtmann and Tally, 2014; Clemmensen et al., 2017). The stomach affects intuition, emotion, and cognition (Mayer, 2011; Klarer et al., 2014, 2018). Such interactions are partly mediated by bidirectional neural signaling along the so-called stomach-brain neuroaxis (Powley et al., 2019; Liu et al., 2018; Kaelberer et al., 2018; Levinthal and Strick, 2020; Browning and Travagli, 2014; Holtmann and Talley, 2014). It includes neural circuits and pathways in both peripheral and central nervous systems (Powley et al., 2019; Mayer et al., 2019; Rebollo et al., 2021).

Prior studies typically characterize the stomach-brain neuroaxis by applying stimulation to the stomach or the brain while recording the resulting brain or gastric responses, respectively. For example, gastric distention can activate the subcortical and cortical networks involved in processing gastric sensation (Ladabaum, 2001; Mayer et al., 2019; Wang et al., 2008; Vandenbergh et al. 2005; Stephan et al., 2003). Gastric electrical stimulation can activate brainstem nuclei and cortical regions and reveal their functional selectivity (Cao et al., 2019; 2021). Likewise, brain stimulation can also modulate gastric motor function (Lyubashina, 2004; Hurley-Gius and Neafsey, 1986). Although these studies offer valuable insights, the stimulation settings used are often artificial, narrowly focused, prone to physiological confounds, and atypical of natural interactions between the stomach and the brain.

Recent studies demonstrate spontaneous interactions between brain activity and gastric electrical activity (Rebollo et al., 2018; Rebollo and Tallon-Baudry, 2021; Choe et al., 2021; Richter et al., 2017; Todd et al., 2021). In their seminal work, Rebollo et al. reports that resting state brain activity observed with functional magnetic resonance imaging (fMRI) is phase locked with gastric slow waves observed with electrogastrography (EGG) (Rebollo et al., 2018). This finding has been replicated by their follow-up study (Rebollo et al., 2021) and an independent group (Choe et al., 2021). In human brains, the stomach-brain synchrony is most pronounced in sensorimotor areas, default-mode network, and cerebellum (Rebollo et al., 2018; Choe et al., 2021) and likely extends further to other networks or systems. In addition, the phase of gastric slow waves is coupled with the amplitude of alpha oscillations in magnetoencephalography (MEG) (Richter et al., 2017) and electroencephalogram (EEG) (Todd et al., 2021), suggesting that the brain activity coupled with gastric activity is of neural origin.

It is also unclear how brain and gut rhythms are synchronized in real-time despite a long distance. The vagus is central to how the brain modulates gastric motor function, such as gastric tone (Azpiroz and Malagelada, 1987), accommodation (Takahashi and Owyang, 1997), motility, and emptying (Lu et al., 2018, 2020), primarily through the dorsal vagal complex in the brainstem or namely the vagovagal reflexes (Powley et al., 2019; Travagli and Anselmi, 2016; Harper et al., 1959; Shapiro and Miselis, 1985; Roger et al., 1995, 1996). The dorsal vagal complex interacts with the hypothalamus, thalamus, amygdala, insular cortex, medial prefrontal cortex, and sensorimotor cortex (Browning and Travagli, 2014; Rinaman and Schwartz, 2004; Han et al., 2018; Levinthal and Strick, 2020; Mayer et al., 2019). Moreover, a subtype of vagal afferent receptors, namely intramuscular arrays (Berthoud and Powley 1992; Brookes et al., 2013), directly innervate interstitial cells of Cajal (ICCs) (Powley et al., 2019). The ICCs initiate the slow wave and pace the stomach (Cajal 1893; Sanders et al., 1996). Taken together, the vagal pathways and their extension in the brain are plausible substrates that bring widespread brain activity into partial synchronization with gastric slow waves.

Here, we hypothesize that the vagal nerves mediate the stomach-brain synchrony in the resting state. We acquired EGG and fMRI simultaneously in rats and evaluated their coherence at the frequency of the gastric slow wave in an attempt to replicate the findings in prior human studies (Rebollo et al., 2018; Choe et al., 2021). Importantly, the use of animal models allowed us to apply bilateral vagotomy and assess its effects on the apparent stomach-brain synchrony. In addition, we also tested the effect of fasting vs. fed (postprandial) states.

## Methods and Materials

### Animals

A total of 18 Sprague-Dawley rats (male, 250-350 g, Envigo RMS, Indianapolis, IN) were used in this study. All animals were housed in a controlled environment with a temperature at 21 ± 1 °C and light on from 6 AM to 6 PM. After diet training and overnight fasting, the animals were divided into three groups for simultaneous acquisition of fMRI and EGG under three different conditions: fed with intact vagal nerves (n=8) or vagotomy (n=5), or fasting with the intact vagus (n=5). All animal procedures followed a protocol approved by the Institutional Animal Care and Use Committee and the Laboratory Animal Program at Purdue University.

### Animal preparation

Through diet training, every animal was preconditioned to be able to voluntarily consume a test meal before the experiment. The test meal was a fixed quantity (5 g) of dietgel (DietGel Recovery, ClearH2O, ME, USA). The test meal had 5.6 kcal, containing 0.03 g protein, 1.16 g carbohydrates, 1 g sugars, 0.055 g dietary fibers, 0.095 g fat, and 0.01g saturated fat. As described elsewhere (Lu et al., 2017), the diet training took about 5 days. In the first two days, 56 g dietgel was placed in the animal cage together with regular chow. In the next three days, the animal was first fasted for 18 hours from 6 PM to 12 PM on the next day and fed only with dietgel (56 g) at 12 PM. The animal was allowed to eat the dietgel from 12 PM to 6 PM; until then it was fasted again for the next day.

After diet training, all animals were fasted overnight. Food was removed from the cage at 6 PM to prepare the animal for the experiment starting at about 12 PM the next day. One group of animals (fasted, n=5) remained fasted during the experiment. Other animals (n=13) were provided a 5 g test meal (dietgel) and were able to quickly consume the meal at the beginning of the experiment. Among those that consumed the test meal, one group of animals had additional vagotomy (n=5) whereas the rest had an intact vagus (n=8). The animals were anesthetized for the vagotomy (if applicable), electrode placement, and fMRI-EGG acquisition in that order.

### Anesthesia

Every animal was initially anesthetized using an induction chamber with 5% isoflurane mixed in oxygen delivered at 1 L/min for up to 5 minutes. Then the animal was transferred to a surgical bench, where 1.5-2.5% isoflurane mixed in oxygen was administered at 1 L/min through a nose cone during surgical vagotomy (for 5 animals after the test meal) and placement of EGG electrodes (for all 18 animals). Then the animal was transferred to a rat holder placed in the MRI scanner and was set up to receive isoflurane through a nose cone and dexdomitor (zoetis, Parsippany-Troy Hills, New Jersey, USA) through subcutaneous infusion. When the animal was set up for MRI, isoflurane was delivered at up to 0.5%, 1 L/min and a bolus of the dexdomitor (12.5 μg/Kg, 0.05 mg/ml) was administered through subcutaneous injection. About 15 minutes later, continuous subcutaneous infusion of the dexdomitor started at 12.5 μg/(Kg x hour) and 0.05 mg/ml, while isoflurane was reduced to 0.1-0.5%. Following that, the infusion rate of the dexdomitor was increased by 12.5 μg / (Kg x hour) for every additional hour to maintain a stable physiological condition over the course of fMRI-EGG acquisition. That is, the heart rate at 250-350 beats per minute, the respiratory rate at 20-60 times per minute, SpO2 above 96%, and the rectal temperature at 36.5-37.5 C°. These physiological parameters were monitored using two systems: Pulse Oximeter and Rat & Mouse Heart Rate Monitor Module (mouseSTAT, Kent scientific, Torrington, CT, USA), and the monitoring and gating system for small animals (model1030, SA instruments, Stony Brook, NY, USA). A heating pad was also controlled to keep the animal warm inside the MRI scanner.

### Vagotomy

Bilateral cervical vagotomy was performed in a group of (n=5) animals after they consumed the test meal. The surgery was performed when the animal was anesthetized with 1.5-2.5% isoflurane as described above. After a toe pinch test, the fur in the cervical area was shaved, followed by scrubbing the skin with 75% ethanol and betadine. Then, a 2-3 cm midline incision was made, starting from the jawline and moving caudally. Skin and soft tissue were separated for the (left or right) cervical vagus nerve to be exposed, isolated and cut. This procedure was performed first for the left side and then for the right side, resulting in bilateral cervical vagotomy. The incision was sutured closed to make the animal ready for attachment of EGG electrodes on the abdominal surface (as described in the next subsection).

### Placement of body-surface electrodes

For all animals, an array of electrodes was attached to the abdominal surface for EGG recordings. The fur on the abdominal skin surface was carefully shaved. We also applied hair remover lotion and waited for 5 min and then removed the extra fur. The shaved skin was further cleaned with sterile saline to prepare for electrode attachment. The electrodes were a 5-by-6 array on a thin 45 x 60 mm^2^ perylene substrate (chronic EMG patch, Microprobes, Gaithersburg, MD, USA). Each electrode in the array was a 1 x 2 mm^2^ platinum contract, with the distance between adjacent electrodes of 7 mm. The array was attached to the skin surface and sutured at four corners and four edges. Additional skin tape was applied to further secure the electrode attachment. In the array, the rows were aligned from left to right and the columns were aligned from superior to inferior (Figure 1A). The intersection of the second row and the third column was on top of the xiphoid. In addition, two adhesive carbon electrodes (Neotech, Valencia, CA, USA) were used as the reference and the ground for EGG, respectively. The reference was about 5 mm superior to the xiphoid on the central line. The ground was on the left hind limb.

**Figure 1.**
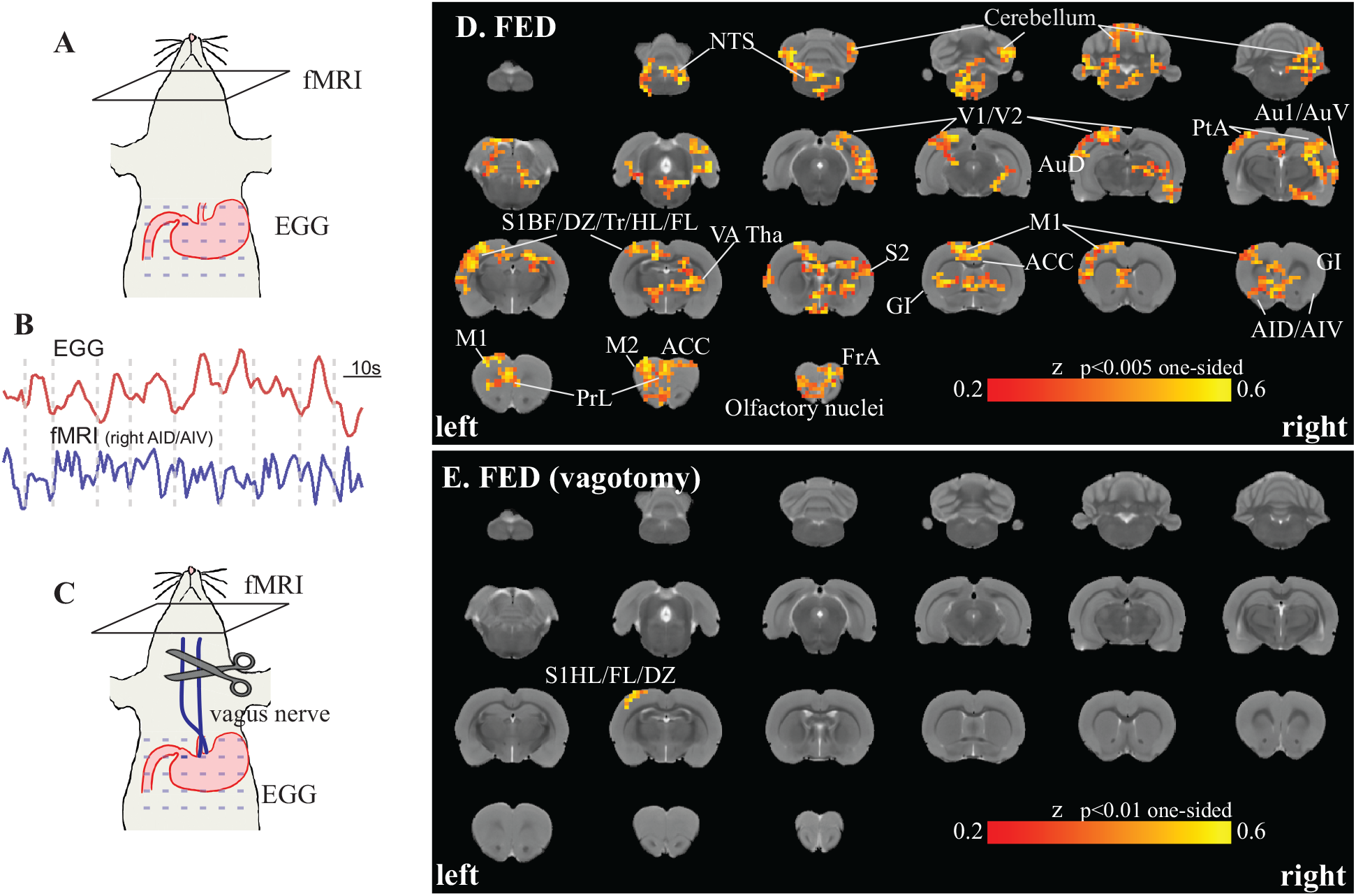
fMRI activity coherent with gastric slow waves. **A** & **C** illustrate concurrent fMRI and EGG in two groups of rats after consuming a test meal with either intact vagus nerves (**A**) or bilateral cervical vagotomy (**C**). **B** shows an example EGG signal collected from xiphoid and the concurrent fMRI signal from one example region (right AID/AIV). The EGG and fMRI signals shown here demonstrate coherence between them at a low frequency around 5 CPM. **D** shows a map of voxels in which fMRI signals were significantly coherent with EGG (one-sided p<0.005, one-way t-test). EGG-coherent cortical and subcortical regions are labeled according to existing rat brain atlases (Paxinos & Watson, 2006; Valdes Hernandez et al., 2011; Papp et al., 2014). **E** shows the EGG-coherent regions in animals with bilateral cervical vagotomy (one-sided p<0.01, one-way t-test). In both **C** and **E**, the underlay shows the T2-weighted images in an existing rat brain MRI template (Valdes Hernandez et al., 2011). The abbreviated labels are explained as the following: ACC, anterior cingulate cortex; AID, agranular insular cortex, dorsal region; AIV, agranular insular cortex, ventral region; FrA, frontal association cortex; GI, granular insular cortex; M1, primary motor cortex; M2, secondary motor cortex; NTS, nucleus tractus solitarius; PrL, prelimbic cortex; PtA, parietal association cortex; S1BF, primary somatosensory cortex, barrel field; S1DZ, primary somatosensory cortex, dysgranular region; S1FL, primary somatosensory cortex, forelimb region; S1HL, primary somatosensory cortex, hindlimb region; S1Tr, primary somatosensory cortex, trunk region; S2, secondary somatosensory cortex; V2, secondary visual cortex; VATha, ventral anterior thalamic nucleus.

### EGG recording

The cutaneous EGG signals were recorded using an electrophysiology system (Tucker Davis Technologies, Alachua, FL, USA) while the animal was placed inside the MRI scanner during concurrent fMRI (see details in the next subsection). The electrodes were connected via Pt/Ir wires and an Omnetics 36 socket nano connector to a headstage (LP32CH-Z-32 Channel Chronic Headstage, Tucker Davis Technologies, Alachua, FL, USA) near the animal. The headstage was connected via a shielded copper cable to an amplifier (PZ5 NeuroDigitizer, Tucker Davis Technologies, Alachua, FL, USA) placed in the MRI room. The amplifier digitized the signals (sampling rate: 24 kHz; bandwidth: DC to 24 kHz; dynamic range: +/-500mV; ADC resolution: 28 bits) and connected to a data acquisition system (RZ2 BioAmp processor, Tucker Davis Technologies, Alachua, FL, USA) outside the MRI room through an optical fiber. The data acquisition system also received and stored a TTL trigger from MRI to synchronize EGG and fMRI acquisition.

### MRI and fMRI acquisition

FMRI was performed with the 7-tesla small-animal MRI system (BioSpec 70/30, Bruker, Billerica, MA, USA) with a volume transmit coil (86 mm inner diameter) and a quadrature surface receive coil. Each animal was placed on a rat holder in the prone position. The head was fixed with two ear bars and a bite bar. The brain’s structure was acquired using a 2-D rapid acquisition with relaxation enhancement sequence with the repetition time (TR) = 5804.6 ms, effective echo time (TE) = 32.5 ms, echo spacing = 10.83 ms, voxel size = 0.125×0.125×0.5 mm^3^, RARE factor = 8, 50 slices, and flip angle (FA) = 90°. FMRI was acquired using a 2-D single-shot gradient-echo (GE) echo-planar imaging (EPI) sequence with TR = 1 s, TE = 16.5 ms, the number of repetition = 1,800, in-plane resolution = 0.5×0.5 mm^2^, slice thickness = 1 mm, 25 slices, FA = 55°, and duration = 30 minutes. Each animal in the fed state (with or without vagotomy) underwent two sessions of fMRI, whereas each animal in the fasted state underwent four sessions of fMRI. The total time of fMRI-EGG acquisition was up to 2 hours in every animal.

### EGG preprocessing

The high sampling rate and broad bandwidth of the EGG recording system allowed for complete separation of MRI-induced artifacts and EGG signals in the frequency domain without any confounding aliasing. Specifically, the signal recorded at each channel was first demeaned and then detrended by regressing out the 5th-order polynomial function that spanned the 30-min duration of each session. A low-pass filter was applied to the detrended signal with the cut-off frequency at 0.45 Hz. The signal was resampled at 1 Hz such that EGG and fMRI shared the same sampling frequency. Although data was recorded from an array of electrodes, analysis in this paper focused on one electrode located at the intersection of the second row and the third column of the array (Figure 1A). Choosing this electrode as the channel of interest was because it was above the antrum of the stomach with peristaltic contractions (Lu et al., 2017). Among all electrodes, the location of this electrode was also most consistent across animals, since it was intentionally positioned to be on top of xiphoid in every animal.

### fMRI preprocessing

The fMRI data was pre-processed using FSL (Jenkinson, et al., 2012), AFNI (Cox, 1996), and in-house software developed in MATLAB. Specifically, we first performed motion correction by registering the fMRI volumes from individual time points to the first volume using *3dvolreg* in AFNI. The first 10 volumes were then discarded to avoid a non-steady state of MR signal. The timing of every slice within each volume was corrected using *slicetimer* in FSL. To align fMRI data to the rat brain template (Valdes Hernandez et al., 2011), fMRI images were aligned to T2w anatomical images from the same animal using *flirt* in FSL, the T2w images were transformed to the template using *fnirt* in FSL, and applying the same transformation to the fMRI images coregistered individual animals’ fMRI data to the same template. The coregistered fMRI signals were further denoised by regressing out the head-motion parameters and a 3rd-order polynomial fit of the slow drift. Lastly, data were spatially smoothed with a 3-D Gaussian kernel with a 0.5-mm full-width at half maximum (FWHM).

### EGG-fMRI coherence

Coherence between EGG and fMRI signals was evaluated at the dominant frequency of gastric slow waves. The slow-wave frequency was identified separately for each animal every 4 minutes to account for its variation across rats and over time. The slow-wave frequency was determined as the frequency with the greatest power spectral density between 0.06 and 0.13 Hz. This frequency range was reasonable since gastric slow waves were expected to be around 5 cpm (or 0.083 Hz) for rats (Tümer et al., 2008). We excluded data from the coherence analysis when the slow-wave frequency was outside the range of 0.06-0.13 Hz.

Coherence between EGG and fMRI signals was evaluated at the slow-wave frequency. It was calculated as the power of the cross-spectral density normalized by the auto-spectral density as expressed in the following equation.

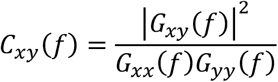

where *x* is the EGG signal, *y* is the voxel-wise fMRI signal, *f* is the slow-wave frequency, *G_xy_*(*f*) is the cross-spectral density, *G_xx_*(*f*) and *G_yy_*(*f*) are auto-spectral density, specifically evaluated at the frequency *f*. The resulting coherence, *C_xy_*(*f*), reflects the power transfer between x and y. In the context of this study, the EGG-fMRI coherence measured the extent to which the gastric slow wave can predict or be predicted by brain activity at a voxel or region via linear systems. It can also account for the potential impact of the hemodynamic impulse response function on fMRI time series.

### Statistical analysis

The voxel-wise coherence was tested for significance by first converting it to a z score for each animal and then applying a one-sample t test to z scores across animals. For each animal, the coherence measured at each voxel was compared against a null distribution, of which the samples were the coherence between randomly phase-shuffled EGG and fMRI signals. Phase shuffling of a signal kept the magnitude but altered the phase of its spectrum. After phase shuffling of EGG and fMRI signals, their coherence was disrupted and reduced to a chance level. The coherence of the original EGG and fMRI signals was compared to the null distribution and converted to a z score by subtracting the mean and dividing by the standard deviation of the chance-level coherence. To test for the significance in the group level, a one-sample t test was voxel-wise applied to z scores obtained from each group of animals (one-sided p<0.01 corrected with a cluster size greater than 10 voxels or 2.5 mm^3^).

The coherence with EGG was summarized with respect to anatomical regions of interest (ROIs). We used 156 ROIs pre-defined according to the rat brain template and atlas. The ROIs included cortical regions (Valdes Hernandez et al., 2011), subcortical regions or nuclei (Papp et al., 2014), and hand-drawn masks for the nucleus tractus solitarius (NTS) and subnuclei in the thalamus (Paxinos and Watson, 2006). For each ROI, the voxel-wise coherence with EGG (in terms of the z score as described above) was first averaged within the ROI for each animal and then tested for significance across animals in the same group using a one-sample t test (with onesided p<0.01 and Bonferroni correction for multiple comparisons). The contrast between different groups, e.g., fed vs. fasted or vagal innervation vs. de-innervation, was tested for significance using two-sample t tests with two-sided p<0.05.

## Results

Prior studies suggested that the human brain had a gastric network, in which regional fMRI activity is phase-locked to the gastric slow wave - an intrinsic gastric rhythm at 0.05 Hz (Rebollo et al., 2018; Choe et al., 2021). Here, we asked whether rat brains had a similar gastric network, and if yes, whether and how this network relied on the vagal nerves that mediate stomach-brain interactions. To address these questions, we simultaneously acquired EGG and fMRI signals in 18 anesthetized rats and evaluated the EGG-fMRI coherence after they voluntarily consumed a 5 g test meal with either an intact vagus (“fed”, n=8) or bilateral cervical vagotomy (“vagotomy”, n=5), or when they remained fasted (“fasted”, n=5). The stomach-brain coherence measured in this way was compared between different conditions (vagal innervation vs. de-innervation and fed vs. fasted) to address whether and how the gastric network relied on intact vagal nerves and varied across gastric states.

### Rat brains also had a gastric network

The slow waves recorded from anesthetized rats after consuming a test meal were 5.36 ± 0.64 (average± standard deviation) CPM or 0.09 ± 0.01 Hz, in contrast to ~3 CPM (or 0.05 Hz) in humans (Chen et al., 1999; Hamilton et al., 1986). We calculated the coherence between the EGG signal from a channel of interest above the antrum (Figure 1A) and the fMRI signal from every voxel in the post-prandial or fed state. See Figure 1B for exemplar EGG and fMRI signals.

In animals with intact vagal nerves, the fMRI signals were coherent with EGG at the slow-wave frequency for 17% brain voxels (one-sided p<0.005, one-sample t test), highlighting a network involving both subcortical and cortical regions (Figure 1C). The brain network, herein referred to as the gastric network, covered the anterior cingulate cortex, insular cortex, medial prefrontal cortex, sensorimotor cortex, auditory and visual cortex, brainstem and cerebellum (Figure 1C). These regions were homologous to the human brain regions phase locked to gastric slow waves (Rebollo et al., 2018; Choe et al., 2021).

The gastric network was asymmetric between the two hemispheres. As shown in Figure 2A, the left hemisphere had more EGG-coherent regions than the right hemisphere. In the sensorimotor cortex, the EGG-fMRI coherence was spreaded across the cortical representations of multiple body parts, whereas the coherence was consistently weaker on the right hemisphere. In contrast, the right insular and auditory cortices were more coherent with EGG than their left counterparts (Figure 2B). The EGG-fMRI coherence tended to be lateralized to the left hemisphere for subcortical regions or nuclei, especially for the cerebellum.

**Figure 2.**
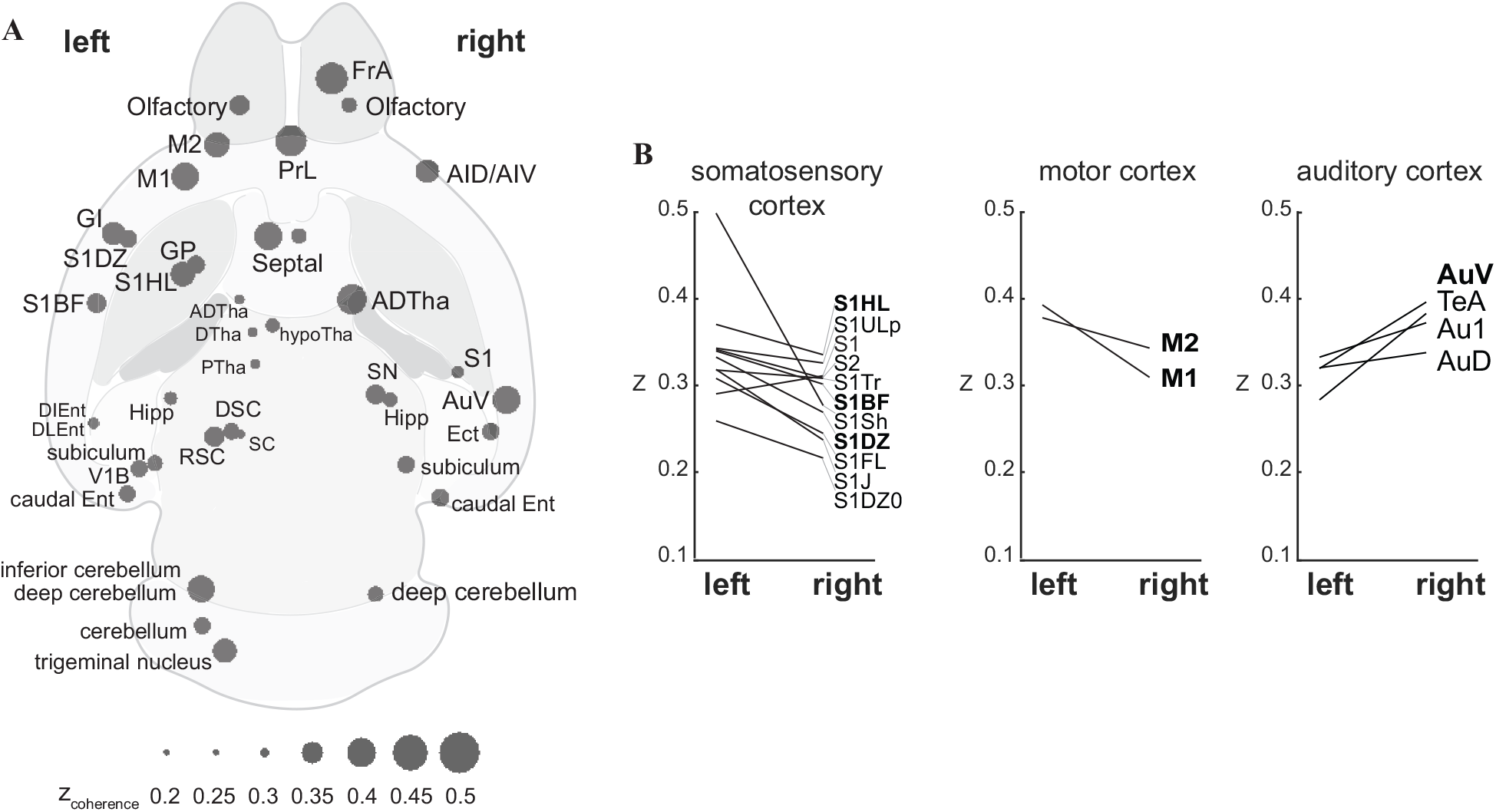
Lateralization of the gastric network. **A** plots the brain regions having significant phase-coupling between the stomach and the brain (one-sample t-test, bonferroni corrected p<0.05). The statistical test was based on the averaged coherence in each region of interest. **B** shows the difference in phase-coupling with EGG between homologous ROIs in the left vs. right hemispheres. Results for the ROIs in the somatosensory cortex, motor cortex, and auditory cortex are shown in three separate subplots shown in the left, middle, and right, respectively. ADTha, anterior dorsal thalamic nucleus; AID, agranular insular cortex, dorsal region; AIV, agranular insular cortex, ventral region; Au1, primary auditory cortex; AuD, secondary auditory cortex, dorsal area; AuV, secondary auditory cortex, ventral area; caudal Ent, caudal entorhinal field; DIEnt, dorsal-intermediate entorhinal field; DLEnt, dorsal-lateral entorhinal field; DSC, deep superior colliculus; DTha, dorsal thalamus; Ect, ectorhinal cortex; FrA, frontal association cortex; FrL, prelimbic cortex, GI, granular insular cortex; GP, globus pallidus; Hipp, hippocampus; hypoTha, hypothalamus; M1, primary motor cortex; M2, secondary motor cortex; Olfactory, olfactory bulb; PTha, posterior thalamus; RSC, retrosplenial cortex; S1, primary somatosensory cortex; S1BF, primary somatosensory cortex, barrel field; S1DZ, primary somatosensory cortex, dysgranular region; S1FL, primary somatosensory cortex, forelimb region; S1HL, primary somatosensory cortex, hindlimb region; S1Sh, primary somatosensory cortex, shoulder region; S1ULp, primary somatosensory cortex, upper lip region; S2, secondary somatosensory cortex; SC, superior colliculus; septal, septal nucleus; SN, substantia nigra; V1B, primary visual cortex, binocular area.

These results suggest that rat brains also have a gastric network that manifests intrinsic coherence between EGG and fMRI signals, consistent with previous findings in human brains.

### EGG-fMRI coherence relied on vagal nerves

We further attempted to identify the peripheral neural pathways underlying the apparent EGG-fMRI coherence. We chose to focus on the vagus nerve, which serves as the primary pathway for rapid and bi-directional neural interactions between the stomach and the brain (Travagli and Anselmi, 2016). Hence, we repeated the experiment in (n=5) animals while applying bilateral cervical vagotomy before EGG-fMRI acquisition (Figure 1D). With bilateral vagotomy, the slow wave frequency was 4.95 ±0.57 CPM, and EGG-fMRI coherence significantly decreased but not completely diminished, while the coherence remained in the primary somatosensory cortex (Figure 1E).

The contrast between the conditions without vs. with bilateral vagotomy was evaluated to characterize the effects of the vagus on EGG-fMRI coherence (Figure 3). In the voxel level, bilateral vagotomy reduced the voxel-wise EGG-fMRI coherence in subcortical areas, such as the nucleus tractus solitarius, pontine reticular nucleus, medial globus pallidus, substantia nigra, thalamus, amygdala, and inferior colliculus, as well as fewer cortical regions in the left dysgranular insular and sensorimotor cortex (Figure 3A). Consistently on the ROI level, bilateral vagotomy significantly decreased the EGG-fMRI coherence (p<0.05, two-sample t test) for most regions in the gastric network (Figure 3B) whereas marginal but non-significant increases were found in fewer regions (Figure 3C).

**Figure 3.**
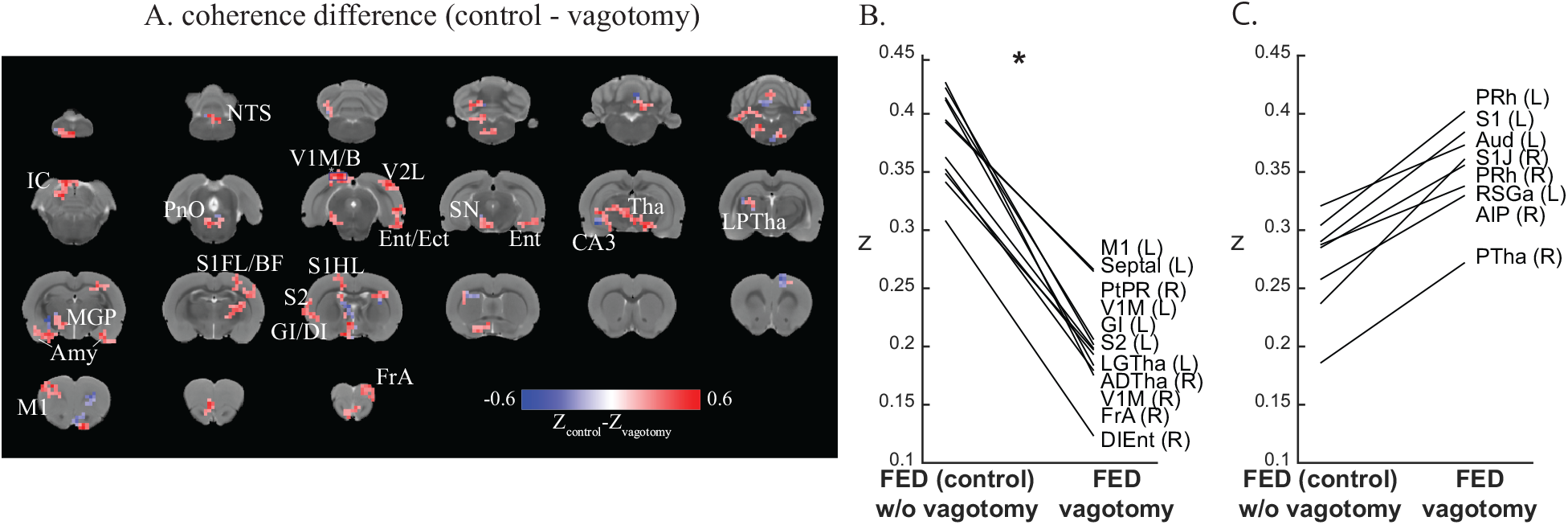
Vagal contributions to postprandial stomach-brain phase-coupling. **A** shows the voxel-level differences of coherence with a threshold of 0.25 between the control group (intact vagal nerves) and vagotomy at the fed state. **B** plots the ROI-level differences in phase coupling (quantified as z scores) for regions where the coupling is stronger with the intact vagus than with vagotomy. The differences shown in this plot are statistically significant (two-way t-test, p<0.05). Similarly, **C** plots the differences for the regions where the coupling is weaker with the intact vagus than with vagotomy, although the differences are not statistically significant. ADTha, anterior dorsal thatlmus; AIP, agranular insular cortex, posterior part; Amy, amygdala; Aud, secondary auditory cortex, dorsal area; CA3, field CA3 of hippocampus; DI, dysgranular insular cortex; DIEnt, dorsal-intermediate entorhinal field; Ect, ectorhinal cortex; Ent, entorhinal cortex; FrA, frontal association cortex; GI, granular insular cortex; IC inferior colliculus; LGTha, dorsal lateral geniculate nucleus; LPTha, lateral posterior thalamus; M1, primary motor cortex; MGP, medial globus pallidus; NTS, nucleus tractus solitarius; PnO, pontine reticular nucleus, oral part; PRh, perirhinal cortex; PTha, posterior thalamus; PtPD, dorsal posterior parietal cortex; PtPR, rostral posterior parietal cortex; RSGa, retrosplenial granular a cortex; S1, primary somatosensory cortex; S1BF, primary somatosensory cortex, barrel field; S1FL, primary somatosensory cortex, forelimb region; S1HL, primary somatosensory cortex, hindlimb region; S1J, primary somatosensory cortex, jaw region; S2, secondary somatosensory cortex; septal, septal nucleus; SN, substantia nigra; Tha, thalamus; V1B, primary visual cortex, binocular area; V1M, primary visual cortex, monocular area; V2L, secondary visual cortex, lateral area.

These results suggest that the vagus is the primary (not necessarily the only) peripheral nerve that mediates the coherence between gastric and brain activities observed with EGG and fMRI respectively.

### EGG-fMRI coherence depended on gastric states

For both rats and humans, the stomach shows continuous and peristaltic contractions in the post-prandial (fed) state and shows irregular and intermittent peristalsis (or migrating motor complex) in the fasted state (Deloose et al., 2012). We asked whether and how EGG-fMRI coherence depended on the fasted vs. fed state. To address this question, we repeated the same experiment in (n=5) animals that remained fasted (after 18-hour fasting) during concurrent fMRI-EGG acquisition. In the fasting state, the slow-wave frequency was 5.02 ± 0.85 CPM. It was comparable with but slightly lower than the slow-wave frequency of 5.36 ± 0.64 CPM in the fed state (Figure 4A).

**Figure 4.**
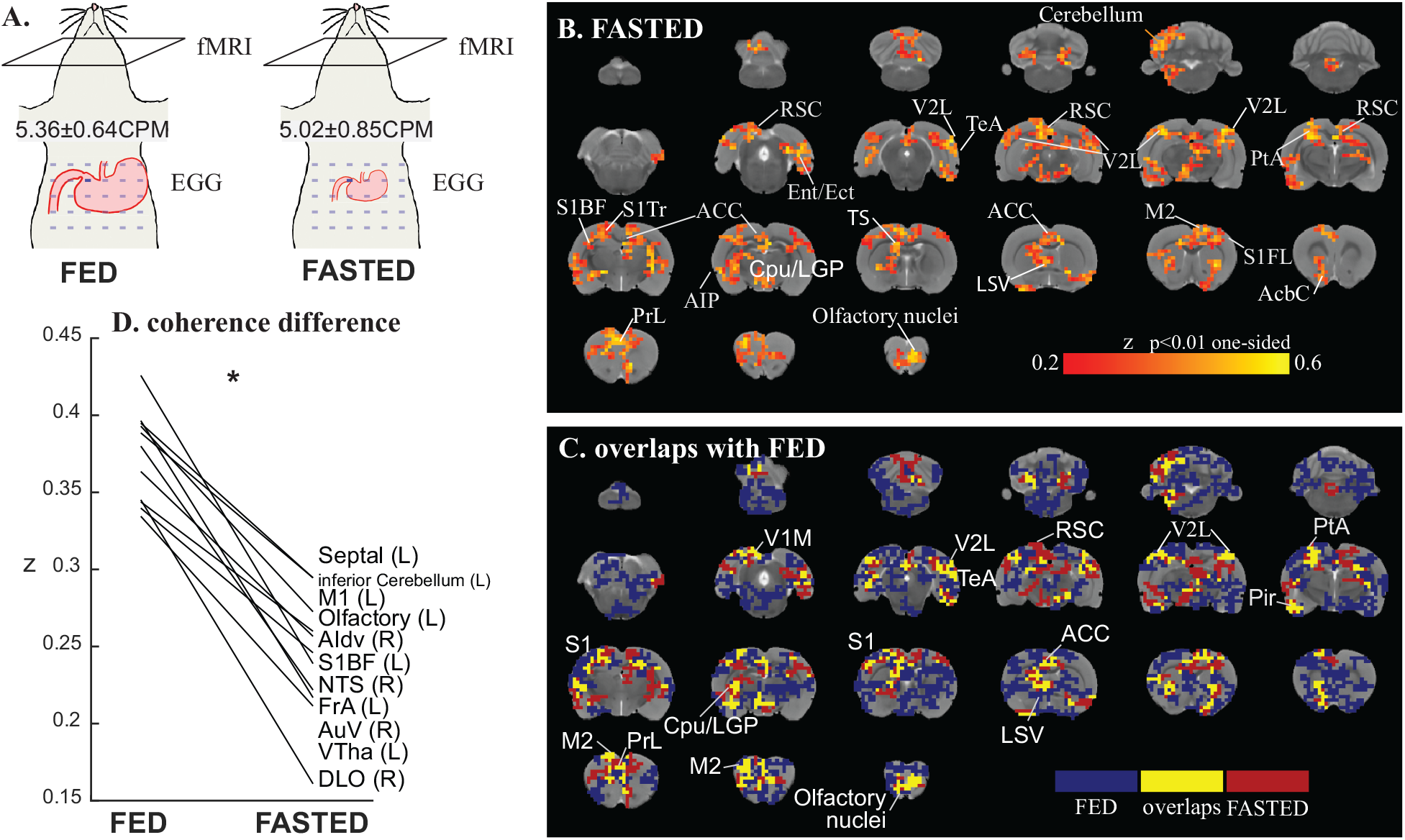
Stomach-brain phase coupling depends on the gastric state. **A** illustrates the fed vs. fasted state and shows the slow-wave frequency (mean±standard deviation) in each state. **B** shows the voxels in which fMRI signals are significantly coherent with EGG in the fasted state (one-sided p<0.01, t-test). **C** shows the overlapping voxels showing significant phase-coupling with EGG in both fasted and fed states with p<0.01. **D** plots the ROI-level differences in phase-coupling between the fed and fasted states, for the regions where such differences are statistically significant (t-test, p<0.05). AcbC, accumbens nucleus, core; ACC, anterior cingulate cortex; AIP, agranular insular cortex, posterior part; AuV, secondary auditory cortex, ventral area; Cpu, caudate putamen (striatum); DLO, dorsolateral orbital cortex; Ect, ectorhinal cortex; Ent, entorhinal cortex; FrA, frontal association cortex. LGP, lateral globus pallidus; LSV, lateral septal nucleus, ventral part; M2, secondary motor cortex; Pir, piriform cortex; PrL, prelimbic cortex; PtA, parietal association cortex; RSC, retrosplenial cortex; S1BF, primary somatosensory cortex, barrel field; S1FL, primary somatosensory cortex, forelimb region; S1Tr, primary somatosensory cortex, trunk region; TeA, temporal association cortex; TS, triangular septal nucleus; V1M, primary visual cortex, monocular area; V2L, secondary visual cortex, lateral area; VTha, ventral thalamus.

We assessed the EGG-fMRI coherence for the fasted state in the same way as for the fed state. In the fasted state, EGG-fMRI coherence was observed at brain regions (Figure 4B) that partially overlapped with those observed in the fed state (Figure 1C). To better appreciate their relationships, Figure 4C shows the intersection and complement of EGG-coherent voxels observed in the fed vs. fasted states. In regions with an overlap between the two states, the coherence with EGG was generally higher in the fed state than in the fasted state (Figure 4D). The ROI-level difference was significant in the primary somatosensory cortex, primary motor cortex, auditory cortex, olfactory nucleus, orbital cortex, septal nucleus, thalamic nuclei, agranular insular cortex, and nucleus of the solitary tract (two-sample t-test, p<0.05).

These results suggest that EGG-fMRI coherence exists in both fasted and fed states but tends to cover complementary subdivisions of a similar set of cortical and subcortical regions.

## Discussion

Here, we present evidence for the coherence between brain activity observed with fMRI and gastric activity observed with EGG in rats. Our findings add to prior observations in humans (Rebollo et al., 2018; Choe et al., 2021) and shed light on an intrinsic gastric network with its hallmark feature being the EGG-fMRI coherence at the gastric pace-making frequency. The substantial reduction in the EGG-fMRI coherence by the bilateral vagotomy supports a primary role of the vagus. The alteration in the gastric network from the fasting to fed state suggests its functional relevance to different gastric conditions.

### Stomach-brain coherence

It is only until recently that the spontaneous interaction between brain activity and gastric activity was studied in humans by simultaneously acquiring EGG along with fMRI (Rebollo et al., 2018; Choe et al., 2021), MEG (Richter et al., 2017), or EEG (Todd et al., 2021). Gastric slow waves were found to be coupled with the amplitude of posterior alpha oscillations (Richter et al., 2017; Todd et al., 2021) or the phase of resting state fMRI activities in sensory and motor cortices (Rebollo et al., 2018; Choe et al., 2020). Since posterior alpha oscillations are correlated with resting-state fMRI signals from sensory and motor cortices and the thalamus (Liu et al., 2012), gastric phase-amplitude coupling with posterior alpha oscillations and phase synchronization with fMRI signals are consistent findings suggesting that the human gastric network has its major components in sensory and motor areas (Rebollo and Tallon-Baudry, 2021).

Our study replicates prior human studies; however, several distinctions should be noted. Coherence used in this study and phase-locking value used in previous studies (Rebollo et al., 2018; Choe et al., 2021) are similar measures of spectral coupling, except that coherence takes into account both amplitude and phase relationships, while the phase-locking value only reflects the phase relationship. Gastric slow waves are of a higher frequency in anesthetized rats (~0.08 Hz) than awake humans (~0.05 Hz), approaching the higher limit of the physiological bandwidth of fMRI. Despite these methodological and physiological differences, the rat gastric network observed in this study covers similar regions as previously reported in humans, including the somatosensory cortex, motor cortex, visual cortex, auditory cortex, cerebellum, and insula cortex. This finding is also consistent with a previous study demonstrating that gastric electrical stimulation can induce both neural and fMRI responses in sensory and motor cortices even at frequencies up to 0.8 Hz (Cao et al., 2019). Together, results from this and prior studies suggest that EGG-fMRI coupling at the gastric slow-wave frequency is an intrinsic phenomenon possibly shared across different species.

Compared to the human gastric network, the EGG-fMRI coherence observed in rats covers more subcortical regions, including the nucleus tractus solitarius and nuclei in the basal ganglia, e.g., the globus pallidus, substantia nigra. These regions reside along the pathways between the autonomic nervous system and the limbic system (Albin et al., 1989; Browning and Travagli, 2014; Parent and Hazrati, 1995) and are likely involved in both autonomic control and emotion regulation (Andresen and Kunze, 1994; Pazo and Belforte, 2002; Pierce and Péron, 2020). In particular, the nucleus tractus solitarius in the brainstem receives vagal afferents that innervate the stomach and relays gastric information to the brain (Browning and Travagli, 2014; Powley et al., 2019).

### Supporting anatomical and functional evidence

The EGG-coherent regions spread across multiple resting state networks in rats (Liang et al., 2011, 2012; Liu et al., 2020). Although the gastric network as a whole is not straightforward to interpret at this stage, most of its constituent regions reported herein have been shown to be structurally connected or functionally associated with the stomach. For example, neural tracing studies have shown that the insular cortex, medial prefrontal cortex, and cerebellum all receive gastric inputs and project to the stomach through polysynaptic connections (up to three synapses) (Browning and Travagli, 2014; Levinthal and Strick, 2020). The somatosensory and motor cortices project to the stomach (up to four synapses) via descending efferent pathways (Levinthal and Strick, 2020).

The involvement of visual and auditory cortex as well as hippocampus in the gastric network is supported by indirect functional evidence from the literature. During slow wave sleep, the firing rate of neurons in the visual cortex was found to be dependent on the phase of duodenal myoelectrical activity (Pigarev et al., 2013) or be responsive to both direct and transcutaneous stimulation of the gut (Pigarev et al., 1994, 2006). The fact that visual thalamus, e.g., lateral geniculate nuclei, receives direct projections from the brainstem, e.g., parabrachial nuclei, suggests a possible pathway for gastric signals to propagate to the visual cortex (Erişir et al., 1997; Uhlrich et al., 1988). Similarly, vagus nerve stimulation at the gastric branch or gastric electrical stimulation was found to modulate neural activity in the auditory cortex (Shetake et al., 2012; Engineer et al., 2015; Cao et al., 2019). Neurons in the hippocampus were also found to be activated by gastric distension or gastric electrical stimulation (Xu et al., 2008, 2009; Wang et al., 2006a, 2006b) likely for sensing satiety and regulating appetite (Davidson et al., 2007, 2009; Kanoski and Grill, 2017).

### Role of the Vagus in the stomach-brain synchrony

Our results suggest that the vagus mediates the apparent stomach-brain synchrony. The neural mechanisms underlying the stomach-brain interaction involve both peripheral and central neural circuits. The peripheral component involves the vagus nerve and the splanchnic nerve. In general, the former is central to regulating gastric motor events (Travagli et al., 2006; Travagli and Anselmi, 2016), and the latter is more involved in visceral pain (Ness and Gebhalt, 1990). Bilateral cervical vagotomy abolishes both vagal afferent and efferent pathways that support vagovagal reflexes. The fact that the vagotomy largely diminished the EGG-fMRI coherence suggests that neural signaling along the vagus nerve is central to maintaining the synchrony between the brain and the stomach. The effects of vagotomy likely extend beyond the dorsal vagal complex, where vagal afferents end and efferents start, through connections between the dorsal vagal complex and other brain regions (Tsurugizawa et al., 2009). Also supporting the role of the vagus is the prior finding that vagus nerve stimulation could activate a similar set of brain regions as the regions highlighted in this study (Cao et al., 2017).

It is likely that intrinsic gastric rhythms are the source that causes brain activity to be synchronized with it. Gastric slow waves observed with EGG reflect an intrinsic rhythm that can be generated solely by the stomach itself. This intrinsic rhythm is thought to originate from and propagate through interstitial cells of Cajal (ICC) (Cajal 1893; Sanders et al., 1996), but likely involves other mechanisms as well (Yin and Chen, 2008; Sarna, 2008). ICCs within the circular and longitudinal layers of the stomach (ICC-IM) are innervated by vagal afferent nerves that branch into intramuscular arrays (Powley and Phillips, 2011; Powley et al., 2019), which serve as the receptors to transmit sensory information to the brainstem (Cao et al., 2021). The vagovagal circuitry in the brainstem may pass the gastric rhythm as the bottom-up input to other regions in the gastric network. Taken together, there is a plausible bottom-up pathway to allow intrinsic gastric rhythms to be passed to the brain and account for the apparent brain-stomach synchrony.

It is possible but less likely that brain activity controls the phase of gastric rhythm in order for the brain to be the cause of stomach-brain synchrony. Given the vagotomy, the rat stomach continued to generate a rhythmic gastric slow wave, despite a slightly lower frequency. Further evidence also suggests that the stomach maintains peristaltic contractions even after the bilateral vagotomy. Although the stomach receives inputs from sensorimotor cortical regions (Levinthal and Strick, 2020), there is no direct evidence suggesting that top-down neural inputs control the phase of the gastric slow wave. The vagal efferent nerves innervate gastric enteric neurons (Powley et al., 2019), which further innervate gastric smooth muscle cells (Furness et al., 2020). It is not clear whether and how the brain controls the ICCs and gastric pace-making activity initiated by the ICCs. However, this does not imply that the brain does not influence EGG. As a gross measure of gastric electrical activity, EGG can result from any synchronized current sources from the stomach, including smooth muscle cells and ICCs. The brain may modulate the excitability of smooth muscle cells and their contributions to the amplitude of EGG, without directly influencing the frequency or phase presumably determined by ICCs. In line with this speculation, a previous study showed that the amplitude of EGG at its dominant frequency varies across different waking-to-sleep stages (Orr et al., 1997). We speculate that the brain senses gastric pace-making activity relayed through vagal afferent nerves, whereas the source of the synchronized rhythms is intrinsic to the gut, as opposed to the brain.

### Gastric mechanical versus electrical activity

It should be noted that body-surface EGG may reflect both gastric pace-making activity and gastric contractions. The gastric pace-making activity is the electrical activity generated by ICCs; it remains largely stationary and continuous across all gastric states. In contrast, gastric contractions vary over time, being strong and continuous after a meal and becoming silent and intermittent in the fasting state (Sarna, 1985; Deloose et al., 2012; Abell 1988; Brandstaeter, et al., 2019; Kim and Malagelada, 1986). The presence of EGG-fMRI coherence in the fasting state suggests that gastric pace-making activity alone may cause the stomach-brain synchrony.

However, gastric contractions may also influence the stomach-brain synchrony, since the pattern of EGG-fMRI coherence was different between the fasting and fed states. However, it is not straightforward to fully disentangle how gastric pace-making activity and contractions cause the stomach-brain synchrony within the scope of this study, because they are often coupled and their coupling are influenced by extrinsic neural control. Addressing this question would ideally require simultaneous electromechanical recordings by placing both mechanical and electrical sensors on the stomach surface during concurrent fMRI. Another way is to attempt to separate electrical slow waves attributed to ICCs and myoelectric activity attributed to muscle cells (Kim and Malagelada, 1986; Sander and Publicover, 1994; Stern et al., 2007).

### Functional significance of the stomach-brain coherence

The functional significance of the stomach-brain coherence is unclear and awaits to be established. Speculatively, this phenomenon is an indication of how the brain monitors the gut in real-time and regulates food intake and digestion (Holtmann and Talley, 2014). It might also indicate how the gut exerts interoceptive influences on brain and behaviors (Critchley and Harrison, 2013; Azzalini et al., 2019). To establish its functional significance, it is necessary to relate the stomach-brain coherence to behavior, such eating behavior, perceptual and cognitive performance. It would also be valuable to evaluate the stomach-brain coherence beyond normal conditions or healthy populations and explore its relationship to various symptoms in diseased conditions.

## Conclusion

In healthy and anesthetized rats, spontaneous brain activity in the “gastric network” is coherent with the intrinsic slow wave from the stomach, consistent with recent findings from humans (Rebollo et al., 2018; Choe et al., 2021). This network involves a number of subcortical and cortical regions that span across the sensory, motor, and limbic systems. Importantly, the vagus nerve is the primary pathway that mediates the stomach-brain coherence.

## Acknowledgement

The authors thank Drs. John Furness, Terry Powley, Leo Cheng, Ulrich Scheven for their valuable discussions. This study is supported by the National Institutes of Health (OD023847, OD030538, AT011665), the University of Michigan, and Purdue University.

